# fastCDS: proteome-scale mapping of protein domains to genomic coordinates

**DOI:** 10.64898/2026.07.18.739381

**Authors:** G. Muñoz-Esquivel, J. I. Fuxman Bass, L. F. Soto-Ugaldi

## Abstract

**Summary:** Mapping protein regions to genomic coordinates underpins the study of exon architecture and the interpretation of clinical variants in their exon context. Existing tools resolve individual queries accurately but scale poorly to proteome-wide analyses. We present fastCDS, a C++ toolkit with command line and Python interfaces for rapid protein-to-genome coordinate mapping from GTF annotations. It matches the accuracy of existing methods while running at least two to three orders of magnitude faster. Mapping all human Pfam domains in seconds, we used the resulting atlas to examine how exonic architecture varies with domain function.

**Availability and Implementation:** fastCDS is freely available under the MIT license at {{https://github.com/SotoLF/fastCDS}} and can be installed with pip install fastCDS or mamba install -c bioconda fastCDS. Pre-built GTF genome indices are archived at Zenodo, DOI: https://zenodo.org/records/21436146.

**Supplementary Information:** Supplementary data are available at Bioinformatics online.

## 1 Introduction

Much of our functional understanding of proteins depends on protein-space annotations, including domains, amino acid motifs, active sites, and variant effects, whereas gene annotations are primarily organized around exon-intron structures. Bridging the two annotation systems requires precise protein-to-genome coordinate mapping. Given a protein identifier and an amino acid range, the corresponding genomic coordinates of all coding sequence (CDS) segments must be determined, accounting for strand orientation and codons that span exon boundaries. This mapping is required to interpret clinical variants in their corresponding exonic context (Landrum *et al*. 2018), visualize protein domains within gene structures, design oligonucleotides targeting a chosen coding region, and study exon shuffling across evolution (Patthy 1999).

The rapid growth of proteome-scale reference resources has turned protein-to-genome mapping into a high-volume task. For example, Pfam annotations (Mistry *et al*. 2021) include more than 48,000 functional domains in human proteins, UniProtKB (UniProt Consortium 2025) reports more than 400,000 sequence features in human proteins, and the AlphaFold Database (Varadi *et al*. 2022) provides residue-level confidence estimates for millions of human protein models. Variant repositories are larger still, containing almost three million ClinVar variants (Landrum *et al*. 2014) with protein-level annotations, and more than six million coding variants in COSMIC (Tate *et al*. 2019).

Despite this need, few dedicated protein-to-genome mapping tools explicitly account for exon structure, and existing approaches are not readily scalable to proteome-wide applications. The Ensembl REST API endpoint (Yates *et al*. 2015) (/map/translation/:id/region) accurately maps protein intervals to genomic coordinates. However, it depends on external web requests and operates at roughly one query per second (q/s), limiting it to small number of regions. Similar mapping can be performed locally using the R proteinToGenome() function in ensembldb (Rainer, Gatto, and Weichenberger 2019), which was later implemented in GenomicFeatures (Lawrence *et al*. 2013). GenomicFeatures offers greater annotation flexibility because it can map against a custom Gene Transfer Format (GTF) file through a TxDb object. However, this flexibility does not improve mapping speed.

Although TransVar (Zhou *et al*. 2015) can perform protein-to-genome coordinate conversion, it was designed for the annotation of variants described using Human Genome Variation Society (HGVS) notation. As a result, when a protein region spans multiple exons, TransVar reports a single genomic span covering the entire region rather than separate intervals for the contributing CDS segments.

A separate group of tools visualizes protein domains in the context of exon-intron structure, including GenePlot (Gonzalez-Ibeas 2022) in Python and the Shiny/R application VisProDom (Wang *et al*. 2022). However, these tools are primarily designed to generate gene-level figures rather than reusable genomic coordinates. Their outputs are therefore difficult to integrate directly into downstream analyses. In addition, neither tool is designed for proteome-scale workflows, which would require hours to days to complete.

Here, we present fastCDS, a self-contained toolkit that makes protein-to-genome mapping practical at proteome scale. fastCDS is distinguished by a unique combination of four capabilities: command-line and Python interfaces, support for arbitrary GTF annotations including custom and non-model genomes, complete transcript reconstruction with publication-quality and interactive visualizations, and an in-memory C++ indexing engine that is at least two to three orders of magnitude faster than existing mapping implementations.

## 2 Implementation and features

fastCDS operates in two stages. First, it reads a GTF file and builds a compact binary index. This index stores, for each protein or transcript, the exon coordinates, CDS intervals, gene identifier, gene name, exon number, and any available MANE select or Ensembl canonical tags. The index is built once per annotation and reused for subsequent mapping jobs. Because the index is built directly from GTF files, fastCDS is not tied to a specific annotation source and supports annotation files from GENCODE (Mudge *et al*. 2025), Ensembl (Dyer *et al*. 2025), and RefSeq-based resources (Goldfarb *et al*. 2025), as well as custom GTF files.

After the index is built, each query is resolved by loading the index into memory, finding the requested protein or transcript, and scanning only the CDS exons for that transcript. fastCDS therefore avoids external web requests, repeated GTF parsing, repeated transcript reconstruction, and the overhead of high-level R or Python objects. Together with the C++ implementation, this makes fastCDS two to three orders of magnitude faster than previous approaches.

For each query region, fastCDS also reconstructs the surrounding transcript structure in one of four modes **(Fig. 1A)**. The coding mode returns the CDS exons and marks the segments that overlap the region, whereas the intron mode reports only the introns falling within its genomic span. The span mode collapses this to the full genomic envelope of the region, while the isoform mode expands it to the complete transcript including 5′ UTRs, CDS exons, introns, and 3′ UTRs. Genomic coordinates are reported in both 0-based BED and 1-based TSV formats, so results can be used directly with bedtools (Quinlan and Hall 2010), IGV (Robinson *et al*. 2011), the UCSC Genome Browser (Kent *et al*. 2002), and DataFrame-based workflows. Any mode can additionally emit a BED12 file with one IGV-ready record per region **(Fig. S1A)**.

**Figure 1.**
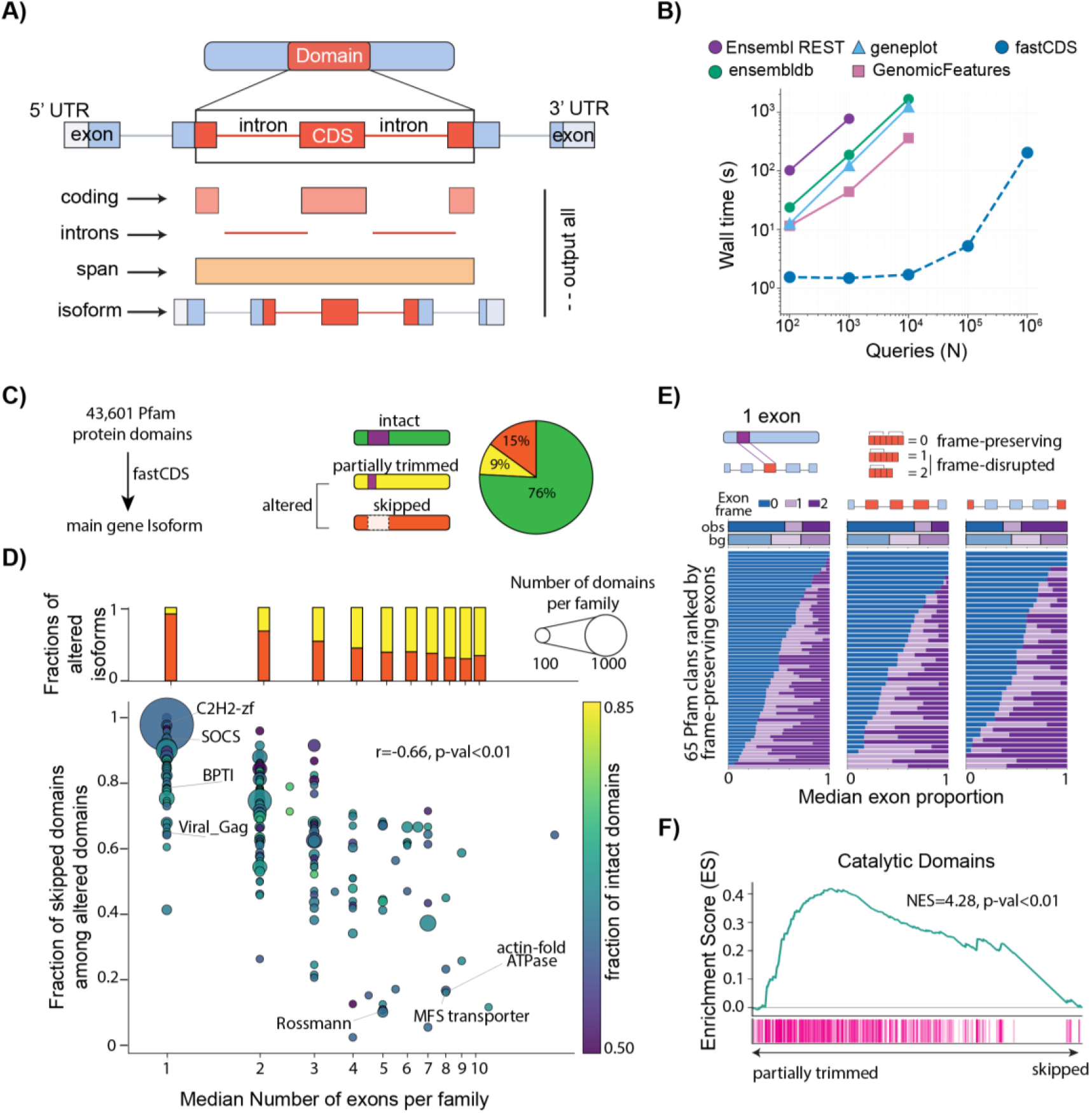
fastCDS enables proteome-scale protein-to-genome coordinate mapping and reveals how exon architecture affects protein domains. **(A)** Schematic of fastCDS coordinate projection. A protein domain is mapped onto the transcript and genomic structure of its coding sequence, producing complementary outputs for coding segments, introns, the full genomic span and complete isoform structure. **(B)** Runtime benchmark across increasing numbers of protein-region queries. fastCDS scales to one million queries, whereas the comparison tools were limited to smaller query sets under the tested conditions. **(C)** Classification of Pfam domains from the main gene isoform as intact, partially trimmed or skipped in alternative isoforms. The pie chart summarizes these outcomes across comparisons involving 43,601 domains. **(D)** Relationship between exon architecture and domain alteration across Pfam families. Points are positioned by the median number of coding exons per domain and the fraction of altered domains classified as skipped. Point size represents the number of domain instances, and color represents the fraction retained intact across isoforms. Stacked bars summarize trimming and skipping by exon count. Spearman’s correlation coefficient and corresponding P value are shown. **(E)** Exon-frame composition of skipped domains. Horizontal stacked bars show the proportions of exons in frame classes 0, 1 and 2 across 65 Pfam clans, ordered by their proportion of frame-preserving exons. Summary bars compare the observed composition with the background distribution. **(F)** Rank-based enrichment of catalytic domains by alteration type. Domains are ranked from partially trimmed to skipped, with vertical marks indicating catalytic-domain instances. The running enrichment score, normalized enrichment score and corresponding P value are shown.

For visualization, fastCDS provides three complementary plotting options from the isoform output. Two Python-based plotting options are available. Using the Matplotlib package, it generates static publication-ready figures in PDF, PNG, or SVG format, including multipage batch plots **(Fig. S1B)**, while with Plotly it generates interactive HTML figures with hover labels and a range slider **(Fig. S1C)**. In addition, a JavaScript option provides a self contained interactive viewer that can be embedded in web pages or Jupyter notebooks, with support for box zoom, a draggable minimap, and switching between true genomic coordinates and intron-compressed layouts **(Fig. S1D)**.

To simplify integration with existing protein annotation workflows, fastCDS also includes converter scripts for common input sources, including InterProScan (Paysan-Lafosse *et al*. 2023), UniProt, and HMMER Pfam (Finn, Clements, and Eddy 2011) outputs. Full output schemas, coordinate conventions, and descriptions are provided in **Supplementary Material 1**.

### 3 Validation and performance

We validated the genomic coordinates produced by fastCDS against four existing tools: the proteinToGenome implementation from ensembldb and GenomicFeatures, TransVar, and the Ensembl REST API. Because these tools depend on specific annotation databases or public API releases, fastCDS was indexed using the matching Ensembl release for each tool. For each comparison, we evaluated 5,000 queries sampled across nine categories designed to include both simple and difficult mapping cases. These included protein regions contained within a single exon, regions spanning multiple exons, codons split across exon boundaries, plus- and minus-strand transcripts, incomplete CDS annotations, selenoproteins, and transcripts with either few or many CDS exons. Full sampling details are provided in **Supplementary Material 2**.

fastCDS reproduced the ensembldb and GenomicFeatures mappings exactly, matching the complete set of genomic intervals for all 5,000 queries. TransVar reports a single enclosing genomic span rather than separate CDS intervals, so direct agreement was evaluated only for queries mapping to a single CDS block.fastCDS matched TransVar exactly for all such queries, while categories potentially containing multiple coding segments were marked as not applicable **(Table S1)**. The Ensembl REST API returned exact matches for 4,953 of 5,000 queries, corresponding to 99.06% overall agreement. Thirty-seven discrepancies occurred, all in the incomplete CDS annotation category, and the other additional 10 queries were not returned by the REST API due to connection errors or API unavailability. For incomplete CDS annotations, the position assigned to residue 1 depends on the mapping convention. fastCDS, ensembldb, and TransVar anchor residue 1 to the first annotated CDS nucleotide, whereas the Ensembl REST API applies the GTF CDS phase, producing an offset of one or two nucleotides. All 37 coordinate differences were within two nucleotides. Additional details are provided in **Supplementary Material 3**.

We next compared runtime and memory usage across five tools: fastCDS, the proteinToGenome() implementations from ensembldb and GenomicFeatures, geneplot, and the Ensembl REST API. To ensure a consistent comparison, all local runtime and memory benchmarks were run using a single thread on one CPU core of a computer equipped with an Intel Core i5 10300H processor and 16 GB of RAM. Using 10,000 queries, fastCDS processed approximately 5,886 q/s, compared with 27.5 q/s for GenomicFeatures and 6 q/s for ensembldb. Thus, fastCDS was more than 200-fold faster than GenomicFeatures and nearly 1,000-fold faster than ensembldb **(Fig. 1B)**. Because of their longer runtimes, geneplot, and the Ensembl REST API were evaluated using 1,000 queries. Geneplot processed approximately 8.1 q/s, and the remotely accessed Ensembl REST API 1.29 q/s **(Fig. S2A)**.

### Alt text

Six-panel figure illustrating the fastCDS workflow, benchmarking performance, and application to protein-domain architecture across isoforms. Panel A shows how a protein domain is projected onto the corresponding transcript and genomic coordinates, producing coding-segment, intron, genomic-span, and full-isoform outputs. Panel B plots runtime against the number of protein-region queries and shows that fastCDS processes substantially larger query sets than the comparison tools, reaching one million queries. Panel C illustrates the classification of domains as intact, partially trimmed, or skipped in alternative isoforms and summarizes the relative frequency of these outcomes. Panel D shows a negative relationship between the number of coding exons spanning a domain and the frequency of complete domain skipping: domains encoded by fewer exons are more often skipped, whereas domains spanning more exons are more often partially trimmed. Panel E compares the proportions of frame-preserving and frame-disrupting exons across 65 Pfam clans and against the genome-wide background. Panel F shows that catalytic domains are concentrated toward the partially trimmed end of a ranking from trimming to skipping.

In addition to its higher throughput, fastCDS used less memory than most of the other tools. Peak memory usage was measured as the maximum resident memory used by each process. Among the tools tested with 10,000 queries, peak memory usage was approximately 808 MB for fastCDS, 1,270 MB for GenomicFeatures, and 1,191 MB for ensembldb.

Loading a prebuilt fastCDS index into memory takes 1.5 seconds. Once loaded, runtime increased linearly with the number of queries. Under these conditions, fastCDS mapped one million queries in 3.4 minutes **(Fig. S2B)**, a scale impractical for the other tools. At their measured throughput, GenomicFeatures and ensembldb would require approximately 10 hours and two days, respectively, for the same workload. For larger analyses, fastCDS also supports parallel processing through OpenMP and allows users to control memory usage by adjusting the batch size **(Fig. S2C)**. Benchmark scripts and the full set of tested parameters are available in the GitHub repository and **Supplementary Material 4**.

## 4 Example application

Previous studies have examined specific relationships between alternative splicing, exon structure, reading-frame preservation and protein domains (Liu and Altman 2003, Magen and Ast 2005). However, to our knowledge, these relationships have not been examined collectively in a quantitative, proteome-wide analysis to identify the general principles governing domain retention among protein isoforms. To address this, we used fastCDS to map all 43,601 human Pfam domains onto the protein-coding isoforms of each gene. Each domain was annotated on the gene’s main isoform and compared with every other protein-coding isoform of the same gene. For each domain-isoform comparison, we recorded whether the domain was fully intact, partially trimmed or skipped altogether in the alternative isoform. Across all comparisons, domains were fully intact in 76% of cases, skipped entirely in 15% and partially trimmed in 9% **(Fig. 1C)**. Domains are therefore generally retained across isoforms, but when altered, they are more often removed as a whole than partially trimmed.

Because isoform differences arise through alternative use of exonic sequence, we evaluated whether exon architecture determines if a domain is skipped completely or partially trimmed. We grouped domains by their number of coding exons and compared these two outcomes across altered isoforms. Single-exon domains, which comprise 36.4% of all domains, were skipped as complete units in 91% of the altered isoforms **(Fig. 1D)**. As the number of coding exons increased, complete domain skipping was less common; the skipped fraction decreased from approximately 54% for domains encoded by three exons to 30-35% for domains encoded by more than ten exons, where partial trimming became the most common outcome. This relationship was also observed across Pfam families, with a strong negative correlation between each family’s median exon count and its skipped fraction (Spearman ρ = −0.66) **(Fig. 1D)**. Compact, single-exon families such as C2H2 zinc fingers, SOCS_box, BPTI and Viral_Gag were usually skipped as complete units. In contrast, multi-exon families such as actin-fold ATPases, MFS transporters and NADP-Rossmann domains were more often partially trimmed.

We next asked whether whole-domain skipping is constrained by the need to preserve the reading frame. Because removal of an internal exon changes the downstream coding sequence unless its coding length is a multiple of three, we compared the frame-preserving properties of internal exons carrying skipped domains with those of internal coding exons genome-wide. Internal exons carrying skipped domains were strongly enriched for frame-preserving exons, with 66% being frame-preserving compared with 42% of internal coding exons genome-wide (OR = 2.75) **(Fig. 1E)**. This bias was specific to internal cassette exons. At transcript ends, where domain loss results from alternative first or last exons and no downstream reading frame is affected, the preference disappeared and was slightly reversed. Finally, we asked whether different patterns of domain alteration were associated with particular functional classes. We ranked altered domains from partially trimmed to completely skipped and tested whether domains assigned catalytic GO terms were concentrated toward either outcome. Catalytic domains were strongly enriched toward the partially trimmed end of the ranking (NES = +4.3) (Fig. 1F). This finding suggests that domain function is associated with how exon architecture is altered across isoforms.

## 5 Conclusion

fastCDS reframes protein-to-genome mapping from a per-query database lookup into a single indexing step, enabling practical application at proteome scale. It can process arbitrary GTFs and support any feature defined by an amino acid range. This flexibility provides a general framework for applications previously infeasible at scale, including exonic annotation atlases, exon-level variant pipelines, and cross-species conservation studies.

## Funding

This work was supported by the National Institutes of Health [grant number R35GM128625 to J.I.F.B.]

## Author contributions

George Munoz Esquivel (Formal analysis [equal], Investigation [equal], Methodology [equal], Software [equal], Validation [equal], Visualization [equal], Writing – original draft [equal], Writing – review & editing), Luis F. Soto-Ugaldi (Conceptualization [equal], Formal analysis [equal], Investigation [equal], Methodology [equal], Software [equal], Supervision [equal], Validation [equal], Writing – original draft [equal], Writing – review & editing), and Juan I. Fuxman Bass (Conceptualization [equal], Formal analysis [equal], Funding acquisition, Investigation [equal], Methodology [equal], Project administration [equal], Resources [equal], Software [equal], Supervision [equal], Validation [equal], Visualization [equal], Writing – original draft [equal], Writing – review & editing)

## Conflicts of interests

None declared.

## Supplementary data

Supplementary data are available at *Bioinformatics* online.

## Data availability

The fastCDS source code, documentation and test data are publicly available from the fastCDS GitHub repository. An archived software release, pre-built genome indices, benchmark query sets, analysis scripts and the processed data underlying the figures are available from Zenodo at DOI: 10.5281/zenodo.21436146. The human gene and transcript annotations used in this study were obtained from Ensembl, and the domain annotations and functional classifications were obtained from Pfam, Ensembl BioMart and InterPro, as described in the Supplementary Methods.

## Notes

https://zenodo.org/records/21436146

### Competing Interest Statement

The authors have declared no competing interest.

