## Supplementary Data for "fastCDS: proteome-scale mapping of protein domains to genomic coordinates"

**Supplementary Materials**

**S1. Input formats, mapping rules, and output files
S1.1 Index construction and query mapping**
The binary index allows each transcript to be retrieved using either its protein or transcript identifier. Protein and transcript identifiers linked to the same transcript therefore produce the same genomic intervals. Version suffixes are removed before lookup, allowing both versioned and unversioned identifiers to be used. RefSeq and custom identifiers are also supported when they are present in the source GTF annotation.

For each query, CDS intervals are processed in translation order. On the plus strand, this corresponds to increasing genomic coordinates, whereas on the minus strand, the CDS intervals are processed in decreasing genomic order. This order is used to assign cumulative CDS positions and map the requested amino acid range back to its contributing genomic intervals. Because the transcript structure is already stored in the index, each query requires only an identifier lookup and traversal of the corresponding CDS intervals. The GTF annotation is not parsed again for later mapping runs.

**S1.2 Query format**

fastCDS accepts a whitespace-separated BED-like input file. Blank lines and lines beginning with # are ignored. Each query can contain the following columns:

1. Identifier: protein or transcript identifier.
2. Amino acid start: 1-based, inclusive.
3. Amino acid end: 1-based, inclusive.
4. Region identifier: optional name carried into the output files.
5. Description: optional free text.

For example:

ENSP00000269305    95    288    P53_DNA_binding

ENSP00000269305    323   356    P53_tetramerization

If no region identifier is provided, fastCDS creates one from the protein identifier and amino acid range. Queries containing only a protein or transcript identifier are processed in structure-only mode. In this mode, fastCDS returns the transcript structure without marking a protein region. Queries that cannot be mapped are written to unmapped_domains.tsv. Reported reasons include an identifier not found in the index, a transcript without CDS annotation, an amino acid range outside the annotated protein length, or a range with no mapped overlap.

**S1.3 Mapping amino acid ranges to genomic coordinates**

fastCDS first joins the CDS intervals of a transcript in translation order and assigns each CDS nucleotide a cumulative position. Amino acid (k) corresponds to CDS nucleotides: 3k-2 to 3k. For an amino acid interval from (a) to (b), the corresponding CDS nucleotide interval is therefore: 3a-2 to 3b. This CDS-relative interval is then projected back onto the genomic CDS segments of the transcript. Using cumulative CDS coordinates allows fastCDS to correctly handle codons whose nucleotides occur on both sides of an exon boundary. In such cases, the genomic coordinates encoding one amino acid are returned as two intervals, one in each CDS exon. For minus-strand transcripts, genomic intervals are still reported with the smaller coordinate first, but their order within the transcript is determined using the strand. The total annotated CDS length is also checked for divisibility by three. Transcripts whose CDS length is not a multiple of three are still processed but are marked using the cds_length_mismatch and cds_nt_remainder fields. The remainder can be zero, one, or two nucleotides. For incomplete CDS annotations, residue 1 is anchored to the first annotated CDS nucleotide in translation order. Differences between this convention and the convention used by the Ensembl REST API are described in Supplementary Methods S3.

**S1.4 Output modes**

The same underlying mapping can be returned through four main output modes.

Coding mode.
Returns the CDS segments of the transcript and identifies the parts that encode the queried protein region. If only part of a CDS exon overlaps the query, the exon is divided into mapped and unmapped portions.

Main files:

- domain_cds_segments.tsv
- domain_cds_segments.bed

Intron mode.
Returns the transcript introns and identifies those located between CDS segments that encode the queried protein region.

Main files:

- domain_introns.tsv
- domain_introns.bed

Span mode.
Returns one continuous genomic interval from the first to the last mapped nucleotide. This interval includes any introns located between the contributing CDS segments.

Main file:

- domain_span_with_introns.bed

Isoform mode.
Returns the full transcript structure, including 5′ UTRs, CDS segments, introns, and 3′ UTRs. Features overlapping the queried protein region are labeled in the output.

Main file:

- isoform_structure.tsv

The all mode generates all of these files together with a run metadata file. A BED12 output can also be added to any mode using the --bed12 option.

**S1.5 Summary and feature tables**

The file domain_mapping_summary.tsv is written for every run and contains one row per query. It reports the main identifiers, chromosome, strand, amino acid range, protein length, genomic span, number of mapped CDS segments, number of CDS exons touched, number of introns crossed, and mapping status.

It also includes:

- MANE Select and Ensembl canonical status;
- whether the full amino acid range was mapped;
- whether the query was processed in structure-only mode;
- whether the CDS length is divisible by three;
- the fraction of the queried region encoded by its largest contributing exon;
- the total length of introns located inside the mapped genomic span.

The CDS, intron, and isoform feature tables use a shared set of columns. These include the query and transcript identifiers, chromosome, strand, feature type, exon number, genomic coordinates, CDS-relative nucleotide coordinates, encoded amino acid positions, overlap status, and the fraction of the queried region contributed by each feature.

Features are numbered in transcript order. For minus-strand transcripts, this differs from genomic coordinate order. When a CDS exon is divided because it is only partly covered by the queried region, all resulting rows retain the same feature identifier and are distinguished using a feature-part number.

**S1.6 Coordinate conventions**

TSV and BED outputs use different coordinate conventions.

- TSV files use 1-based, inclusive coordinates, matching the source GTF.
- BED files use 0-based, half-open coordinates, following the BED standard.

For example, a GTF interval from 7,676,219 to 7,676,272 is written in BED format as 7,676,218 to 7,676,272. Both intervals represent the same 54 nucleotides. Interval length is calculated as: end - start + 1 for TSV coordinates, and: end - start for BED coordinates.Fields that do not apply to a row are reported as NA.

**S1.7 BED and BED12 outputs**

The standard BED outputs contain six columns: chromosome, start, end, name, score, and strand.

The BED files include:

- mapped CDS segments;
- introns inside the mapped genomic span;
- one continuous span covering the full mapped region.

The optional domain_blocks.bed file uses BED12 format and contains one record per query. Each block represents one mapped CDS segment, allowing regions encoded by several exons to be displayed as a single track item in IGV or the UCSC Genome Browser. For structure-only queries, no mapped-region BED records are produced.

**S1.8 Input converters**

fastCDS includes converters for InterProScan, UniProt, and HMMER/Pfam outputs. Each converter extracts the protein identifier, amino acid start and end coordinates, annotation identifier, and optional description, and writes them in the BED-like format accepted by fastCDS map.

The protein identifier in the converted file must match a protein or transcript identifier present in the fastCDS GTF index. Inputs already labeled with Ensembl protein identifiers can therefore be used directly. Inputs labeled with UniProt accessions must first be linked to Ensembl protein identifiers using either Ensembl cross-references contained in the UniProt record or a release-specific UniProt–Ensembl cross-reference table.

The InterProScan converter reads standard TSV output generated with interproscan -f TSV. The protein identifier is taken from the first column and therefore matches the identifier used in the input FASTA. The converter extracts the amino acid start and end positions, signature accession, analysis type, and available InterPro annotation. If the FASTA was labeled with Ensembl protein identifiers, the converted file can be passed directly to fastCDS map. If UniProt accessions were used, an Ensembl cross-reference table is required.

The UniProt converter accepts UniProt flat files or REST JSON records and extracts annotated protein features and their amino acid coordinates. Ensembl protein identifiers are obtained from the cross-references included in the UniProt record or from a release-specific cross-reference table. If one UniProt accession maps to several Ensembl proteins, one query is generated for each matching protein.

The HMMER converter reads per-domain output generated with --domtblout from hmmscan or hmmsearch. The protein identifier is inherited from the input FASTA, while the protein alignment start and end are used as the query coordinates. In hmmscan output, the protein identifier is read from the query field; in hmmsearch output, it is read from the target field. Pfam model coordinates are not used because fastCDS requires coordinates on the protein sequence. As with InterProScan, outputs labeled with Ensembl protein identifiers can be used directly, whereas UniProt-labeled outputs require a corresponding UniProt–Ensembl mapping.

All converters produce a file containing the Ensembl protein or transcript identifier, amino acid start, amino acid end, annotation identifier, and optional description. This file can then be passed directly to fastCDS map.

**S2. Validation set construction**

Mapping accuracy was tested using a fixed set of 5,000 protein-region queries. Each query contained a protein identifier, a starting amino acid, and an ending amino acid. The expected output was the set of genomic intervals encoding that protein region. The query set was generated from the human GRCh38 Ensembl release 86 GTF annotation. Only transcripts annotated as protein coding and containing a protein identifier were included. Transcripts with fewer than nine CDS nucleotides were excluded. CDS exons were ordered according to the direction of translation, from the beginning to the end of the coding sequence. This corresponds to increasing genomic coordinates on the plus strand and decreasing genomic coordinates on the minus strand.

For each transcript, we recorded the strand, number of CDS exons, protein length, gene symbol, and whether the CDS was annotated as incomplete. A transcript was considered incomplete when it contained a cds_start_NF or cds_end_NF tag. Protein length was calculated by dividing the total CDS length by three. We also identified amino acids encoded by codons split across exon boundaries. A codon was considered split when the cumulative CDS length at an exon boundary was not divisible by three.

Queries were sampled using a fixed random seed. Rather than sampling all queries without restriction, we divided them among nine categories to include both common and difficult mapping cases. Sampling was performed with replacement, meaning that the same transcript could be selected more than once with different amino acid ranges. The category filters were not mutually exclusive, so a transcript could be eligible for more than one category. However, each query was assigned only to the category from which it was sampled.

The nine categories were generated as follows:

**Single-exon protein regions (n = 1,000).** One CDS exon containing at least two complete codons was selected, and an amino acid range located entirely within that exon was sampled. These queries were expected to map to one continuous genomic interval.

**Multi-exon protein regions (n = 1,000).** Transcripts with at least two CDS exons were selected. The amino acid range began in one CDS exon and ended in a later CDS exon, ensuring that the region crossed at least one exon boundary.

**Codons split across exon boundaries (n = 500).** Transcripts containing at least one codon split between two CDS exons were selected. A short amino acid range containing the split codon was then sampled. These queries tested whether the nucleotides encoding a single amino acid were correctly divided between two genomic intervals.

**Plus-strand transcripts (n = 1,000).** A random amino acid range was sampled from a transcript on the plus strand.

**Minus-strand transcripts (n = 1,000).** A random amino acid range was sampled from a transcript on the minus strand. These queries tested whether CDS exons were processed in translation order rather than genomic coordinate order.

**Incomplete CDS annotations (n = 200).** A random amino acid range was sampled from a transcript carrying a cds_start_NF or cds_end_NF tag. These queries tested how each tool handled transcripts whose coding sequence was incomplete at the 5′ or 3′ end.

**Selenoproteins (n = 100).** A random amino acid range was sampled from transcripts belonging to a curated set of 25 human selenoprotein genes. These genes contain an in-frame UGA codon that encodes selenocysteine rather than terminating translation. This category tested whether amino acid numbering continued correctly after the UGA codon.

**Single-CDS-exon transcripts (n = 100).** A random amino acid range was sampled from a transcript containing exactly one CDS exon.

**Transcripts with many CDS exons (n = 100).** A random amino acid range was sampled from a transcript containing more than 20 CDS exons. These queries tested mapping across transcripts with highly divided coding regions.

The same 5,000 protein identifiers and amino acid ranges were used in each tool comparison. Because each comparison tool supports a different Ensembl release, fastCDS was indexed separately using the release supported by that tool. fastCDS was compared with ensembldb using Ensembl v86, with TransVar using Ensembl v95, and with the Ensembl REST API using Ensembl v115.

For the ensembldb and Ensembl REST API comparisons, the genomic intervals returned for each query were sorted and compared as sets of chromosome, start, and end coordinates. Chromosome names were standardized before comparison so that names such as 1 and chr1 were treated as equivalent.

TransVar reports one genomic span covering the full protein region rather than separate intervals for each contributing CDS exon. For this comparison, the fastCDS intervals were therefore reduced to their outer genomic boundaries before comparison. Exact agreement required both tools to report the same chromosome and the same outer start and end coordinates.

Agreement was calculated across the full query set and separately for each sampling category. Queries were also divided according to whether the transcript had a complete or incomplete CDS annotation. This second grouping was used to examine the differences observed between fastCDS and the Ensembl REST API, as described in Supplementary Methods S3.

**S3. Comparator setup and mapping differences**

**S3.1 Comparator configuration**

Mapping accuracy was evaluated against GenomicFeatures::proteinToGenome(), ensembldb::proteinToGenome(), TransVar, and the Ensembl REST API. For ensembldb, proteinToGenome() was run using EnsDb.Hsapiens.v86. GenomicFeatures was run using CDS intervals grouped by transcript with cdsBy(edb, by = "tx"); the resulting GRangesList was generated once and retained in memory during the benchmark. TransVar was run with:

transvar panno -l <query_file> --ensembl
using its Ensembl v95 database.

Ensembl REST queries were submitted to: /map/translation/{id}/{start}..{end}
using Ensembl release v115.

The matching Ensembl release was used to build the fastCDS index for each comparison. Query construction and interval comparison are described in Supplementary Material S2.

**S3.2 Selection of the ensembldb database**

The ensembldb proteinToGenome() function requires an EnsDb database containing links between protein identifiers, transcripts, and CDS intervals. EnsDb databases created directly from newer GTF files with ensDbFromGtf() did not contain the protein information required by the function.

We also tested a human EnsDb database for Ensembl release 113 obtained from AnnotationHub. In this database, proteinToGenome() returned “No CDS found” for 2,736 of the 5,000 queries because many proteins were not linked to CDS intervals. We therefore used EnsDb.Hsapiens.v86, the most recent prebuilt human EnsDb tested that completed the full query set.

**S3.3 TransVar database coverage**

The TransVar database contained approximately 0.1% fewer transcripts than the Ensembl v95 GTF from which it was built. A small number of transcripts present in the GTF were therefore unavailable in TransVar.

To prevent missing database entries from being counted as coordinate differences, the TransVar query set was sampled only from transcripts present in both the Ensembl v95 GTF and the TransVar database.

**S3.4 Incomplete CDS annotations**The differences between fastCDS and the Ensembl REST API were restricted to transcripts annotated with cds_start_NF or cds_end_NF. In these transcripts, the annotated CDS does not contain its complete 5′ or 3′ end, and the first available CDS segment may begin within a partial codon.

fastCDS treats the first annotated CDS nucleotide in translation order as the start of residue 1. ensembldb and TransVar follow the same convention. The Ensembl REST API instead applies the CDS phase stored in the annotation. The phase indicates whether zero, one, or two nucleotides must be passed before the next complete codon begins.

For a phase of one or two, the REST mapping is shifted relative to the first annotated CDS nucleotide. In protein regions spanning several exons, this initial shift can also change how the mapped interval is divided at later exon boundaries. Because fastCDS and the REST API were evaluated using the same Ensembl release, these differences reflect the convention used for incomplete CDS annotations rather than differences between annotation versions.

**S4. Performance methodology**

**S4.1 Benchmark environment and measurement**

Comparative runtime, throughput, and memory benchmarks were performed on a laptop equipped with an Intel Core i5-10300H processor and 16 GB of RAM. All local tools were restricted to a single CPU core and were run sequentially to avoid competition for computational resources. The fastCDS thread and batch-size benchmarks were performed separately on a workstation equipped with an Intel Core i9-14900K processor and 128 GB of RAM.

Each tool was launched as a separate process using a common benchmarking script. Total wall time was measured from process launch until the program terminated after writing all output files. This measurement therefore included loading the required index, database, or annotation object, query processing, and output generation. Peak resident memory was recorded using os.wait4 and its ru_maxrss value. Reported values represent the median across three replicate runs.

**S4.2 Comparative runtime, throughput, and memory**

Runtime scaling was evaluated using query sets of increasing size, ranging from 10^2^ to 10^6^ protein intervals when practical for the tool being tested. fastCDS, GenomicFeatures, and ensembldb were evaluated using workloads of up to 10,000 queries for the direct throughput comparison. Because geneplot, and the Ensembl REST API required substantially longer runtimes, their throughput was evaluated using 1,000 queries.

Throughput was calculated as the number of completed queries divided by the total wall time. Peak resident memory was measured during the same benchmark runs. For the Ensembl REST API, the reported memory usage corresponds only to the local client process and is not directly comparable with that of the local tools because the coordinate mapping is performed on the Ensembl server.

**S4.3 Parallel scaling and batch-size effects**

fastCDS parallelizes independent queries using OpenMP, with the number of threads controlled by the --threads option. Results are written in the original input order, so changing the number of threads does not affect the output.

The --batch-size option controls the number of completed queries retained in memory before their results are written to disk. After each batch is written, its buffered records are released before the next batch is processed. Batch size therefore affects memory usage without changing the mapped coordinates.

The effects of thread number and batch size were evaluated using a fixed workload of one million queries. Thread counts of 1, 2, 4, 8, and 16 were tested with batch sizes of 10,000, 50,000, and 100,000 queries, as well as a one-shot configuration in which all one million queries were processed as a single batch. Wall time and peak resident memory were recorded for each combination.

**S5. Pfam atlas & cross-isoform domain behaviour**

**S5.1 Domain architecture on the main isoform**
Human Pfam-A domain instances on the canonical isoform of each gene were obtained from Ensembl BioMart (release 115) as (ENSP, Pfam accession, aa_start, aa_end) records and mapped to genomic coding intervals with fastCDS map --output coding. The analysis set comprised domains lying on the main isoform of genes with ≥ 2 protein-coding isoforms; domains in single-isoform genes cannot be compared across isoforms and were excluded, leaving 43,601 domains. For each domain, n_cds_exons was defined as the number of distinct CDS exons spanned (1 denoting a single-exon domain). Each Pfam accession was assigned to its Pfam clan using Pfam-A.clans.

**S5.2 Isoform universe and cross-isoform projection**
The release-115 GTF was parsed to one row per protein-coding isoform, retaining genes with ≥ 2 coding isoforms. Each gene's reference ("main") isoform was taken as its MANE Select transcript when present, otherwise the Ensembl canonical transcript, otherwise the longest CDS. Domains on the main isoform were projected to their genomic coding intervals (fastCDS map --output coding), and every other isoform of the same gene was mapped structure-only (--output isoform). For each (domain, target-isoform) pair, coverage was computed as the domain coding bp retained as CDS in the target divided by the domain coding bp, and the pair was classified as fully intact (domain retained whole and in frame), skipped (coverage < 0.15), or partially trimmed (0.15 ≤ coverage < 1).  Pooled across all 505,361 domain-target-isoform comparisons — every one of the 43,601 reference-domain instances projected onto each remaining coding isoform of its gene — 75.8 % of comparisons were fully intact, 15.0 % skipped and 9.2 % partially trimmed. These per-comparison shares are pooled over all comparisons rather than averaged per domain, and are computed on the full 43,601-domain set, not the ≥ 3-pair subset defined in S5.3.

**S5.3 Per-domain intactness**
Each domain was summarized by its intactness, the fraction of its gene's isoform pairs in which it was fully intact. Among a domain's altered (non-intact) pairs, the skipped fraction was the number of skipped pairs divided by all altered pairs, with the complementary quantity defining the trimmed fraction. The intactness analysis set comprised the 27,928 domains testable in ≥ 3 isoform pairs (3,932 Pfam families across 545 clans in 10,562 genes). Two filters were applied in the subsequent analyses: a domain was mode-callable when it had ≥ 3 altered pairs, and variable when 0.10 ≤ intactness ≤ 0.90 (excluding near-constitutive and near-always-absent domains).

**S5.4 Mode of alteration by exon architecture**
Variable domains were grouped by exact n_cds_exons (capped at 10+), and for each bin the mean per-domain skipped fraction and mean trimmed fraction were computed. Separately, variable domains were aggregated to the Pfam-family level for families with ≥ 15 variable domains (n = 192); for each family the median n_cds_exons and the mean skipped fraction were computed, and their association across families was assessed by Spearman rank correlation.

**S5.5 Reading-frame preservation of skipped exons**
All CDS exons were extracted from the release-115 GTF, and each was assigned a coding length (end − start + 1) and a phase class equal to that length modulo 3, where 0 denotes a frame-preserving exon (its removal deletes whole codons without shifting the downstream frame) and 1 or 2 a frame-disrupting exon. Within each protein, CDS exons were ordered in translation direction; the first and last were labelled terminal and the remainder internal. Each domain was assigned to its carrying CDS exon by midpoint containment. The analysis was restricted to exons carrying a cleanly skipped single-exon domain (single-exon with skipped fraction ≥ 0.8), deduplicated to unique (clan, exon) pairs, and split by carrying-exon position (all, internal, terminal). For each clan the phase composition (fractions frame-preserving, mod 1, mod 2) was computed and ranked by frame-preserving fraction, retaining clans with ≥ 6 exons for the all-position analysis and ≥ 5 for the internal and terminal analyses. For each position class, the pooled phase composition of skipped-carrying exons was compared with the genome-wide CDS-exon background by two-sided Fisher exact test.

**S5.6 Enrichment of catalytic domains at the trimmed end**
Variable, mode-callable domains (n = 11,749; single- and multi-exon combined) were ranked in descending order by the signed mode metric (trimmed fraction − 0.5). Catalytic domains were defined as instances of Pfam families mapping, via InterPro pfam2go, to GO:0003824 (catalytic activity) or any descendant term, propagated to the instance level (1,065 catalytic domains in the ranked list). Enrichment was tested by pre-ranked, weighted gene-set enrichment analysis, with per-domain weight equal to the absolute ranking metric and the enrichment score defined as the maximum deviation of the weighted running-sum statistic. Significance was assessed from 10,000 gene-set-label permutations: the normalized enrichment score was the enrichment score divided by the mean absolute enrichment score over same-sign permutations, and a one-sided empirical p-value was computed as the fraction of permutations at least as extreme in the observed direction, capped at a minimum of 1/10,000.

**Supplementary Figures**

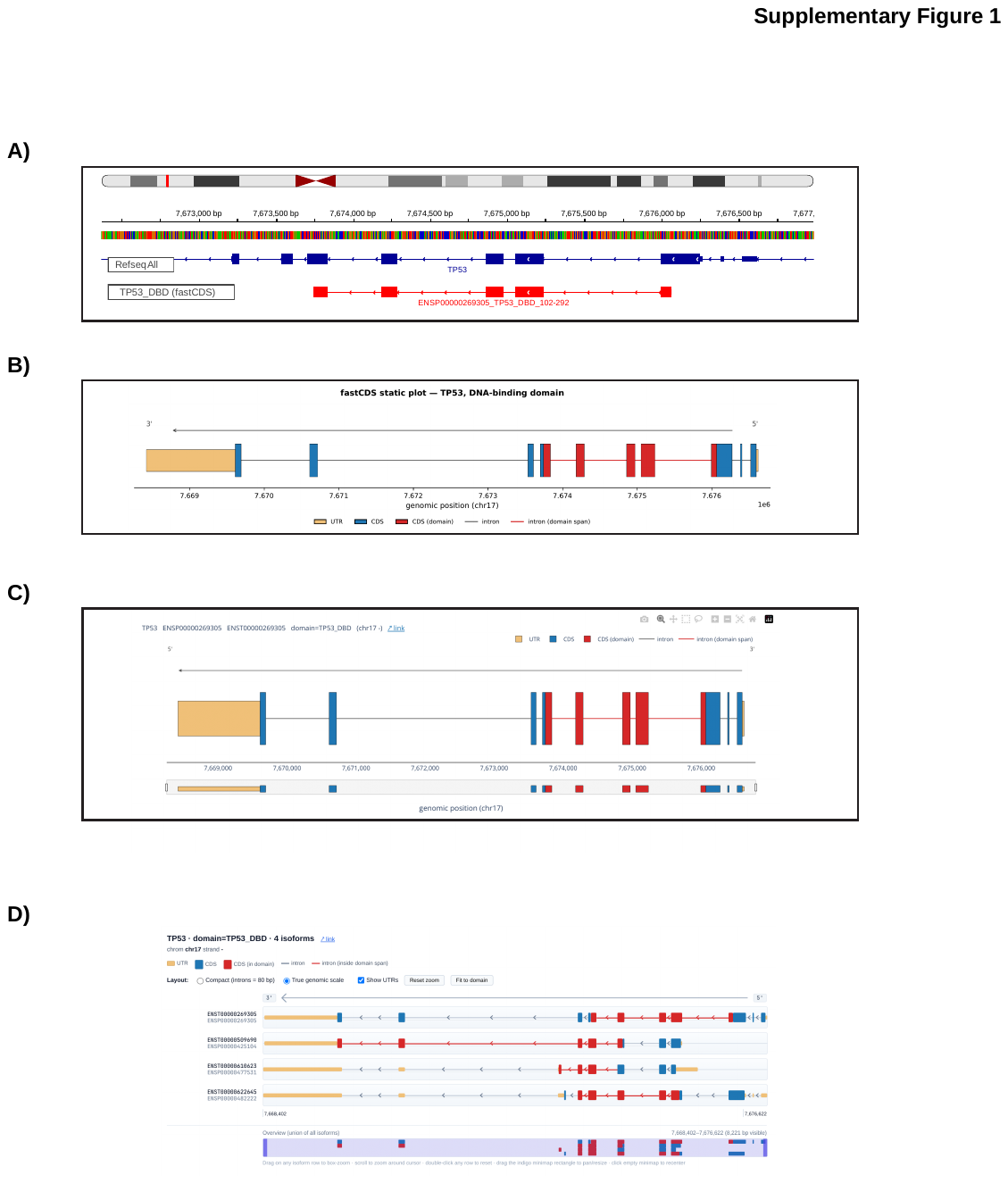

**Figure S1. fastCDS output formats and visualization options.
A)** IGV view of a fastCDS BED12 export. fastCDS writes each queried protein region as a single BED12 record. Coding segments are displayed as connected (red) blocks in IGV or other genome browsers. The example shows a TP53 protein region aligned to the RefSeq gene model, with BED12 blocks corresponding to the coding segments that encode the queried protein feature.

**(B)** Static Matplotlib visualization of the same TP53 protein region, showing the complete transcript structure and the genomic segments that encode the domain.

**(C)** Interactive Plotly HTML visualization, including hover information, zoom controls, and a range slider for navigation along the genomic locus.

**(D)** JavaScript viewer showing the mapped TP53 domain across four protein-coding isoforms. The viewer supports comparison of isoform structures, display at true genomic scale or with compressed introns, optional UTR display, and navigation through a shared overview panel. In panels **B–D**, UTRs are shown in yellow, CDS segments outside the queried region in blue, CDS segments encoding the queried region in red, introns in gray, and introns within the genomic span of the queried region in red.

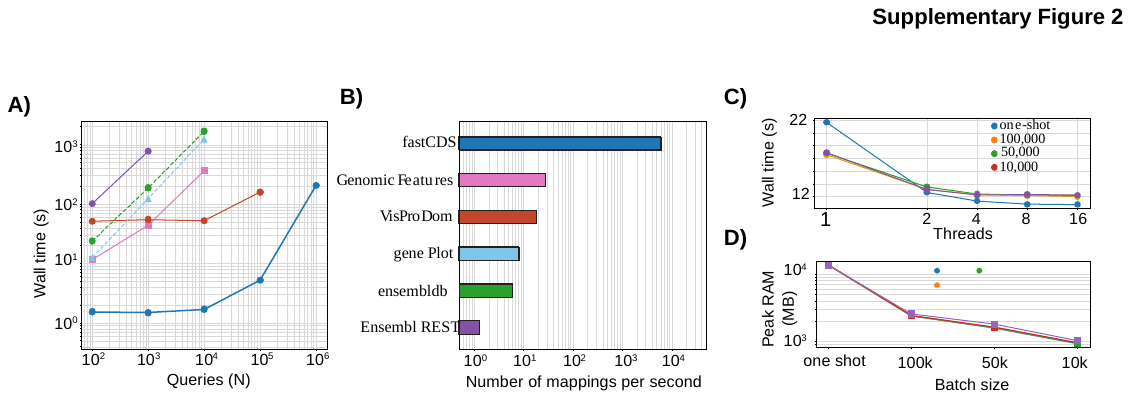

**Figure S2. Runtime scaling, throughput, and the effects of parallelization and batch size.**

**(A)** Log line plot showing wall time as a function of the number of protein-region queries for fastCDS and the comparison tools, with each tool shown in a different color.

**(B)** Horizontal bar plot summarizing the corresponding mapping throughput, measured as queries processed per second. Throughput for fastCDS, GenomicFeatures, and ensembldb was calculated from the 10,000-query benchmark, whereas geneplot, and the Ensembl REST API were evaluated using 1,000 queries.

**(C)** Line plot showing the effect of increasing thread number on the time required to process one million queries at different batch sizes.

**(D)** Line plot showing peak RAM usage for the same one-million-query analysis across batch sizes and thread numbers.

**Supplementary Tables**

**Table S1. Agreement with fastCDS by tool and query category**

Exact coordinate agreement with fastCDS across nine query categories (5,000 protein-region queries total), with each comparator evaluated on its matching Ensembl release (ensembldb and GenomicFeatures, v86; TransVar, v95; REST API, v115) and fastCDS re-indexed accordingly. REST off, coordinate difference ≤2 nt; REST no map, no mapping returned by the REST API; NA, not applicable.

| **Category** | **n** | **ensembldb%** | **Genomic Features%** | **TransVar%^a^** | **REST exact** | **REST off** | **REST no map** | **REST%** |
| --- | --- | --- | --- | --- | --- | --- | --- | --- |
| single_exon_domain | 1,000 | 100 | 100 | 100 | 998 | 0 | 2 | 99.8 |
| multi_exon_domain | 1,000 | 100 | 100 | NA | 999 | 0 | 1 | 99.9 |
| codon_split_boundary | 500 | 100 | 100 | NA | 498 | 0 | 2 | 99.6 |
| plus_strand_gene | 1,000 | 100 | 100 | NA | 997 | 0 | 3 | 99.7 |
| minus_strand_gene | 1,000 | 100 | 100 | NA | 1,000 | 0 | 0 | 100.0 |
| cds_incomplete | 200 | 100 | 100 | NA | 163 | 37 | 0 | 81.5 |
| selenoprotein | 100 | 100 | 100 | NA | 98 | 0 | 2 | 98.0 |
| single_exon_gene | 100 | 100 | 100 | 100 | 100 | 0 | 0 | 100.0 |
| many_exon_gene | 100 | 100 | 100 | NA | 100 | 0 | 0 | 100.0 |
| **OVERALL** | **5,000** | **100** | **100** | **NA** | **4,953** | **37** | **10** | **99.06** |

*a. TransVar returns one genomic span per query and does not report the individual contributing CDS intervals, so it was evaluated only for the single-exon protein-region and single-CDS-exon transcript categories.*

**Table S2. Mapping speed and peak memory usage**

Single-core wall-time and peak-memory benchmarks (Intel Core i5-10300H, 16 GB RAM), measured from program launch until all results were written and including index, database, or annotation loading. The Ensembl REST API memory value reflects only the local client, since mapping is performed on the Ensembl server.

| **Tool** | **Queries s⁻¹** | **Peak RSS (MB)** | **N** |
| --- | --- | --- | --- |
| fastCDS | 5,886 | 808 | 10,000 |
| GenomicFeatures | 27.5 | 1,270 | 10,000 |
| geneplot | 8.1 | 317 | 1,000 |
| ensembldb | 6.0 | 1,191 | 10,000 |
| Ensembl REST | 1.29 | 28 | 1,000 |
